# Exploring the multi-protein assembly of the enzymes of the *de novo* purine nucleotide biosynthetic pathway from *Pseudomonas aeruginosa*

**DOI:** 10.1101/2025.07.15.664858

**Authors:** Nour Ayoub, Nicolas Pietrancosta, Quentin Giai Gianetto, Gouzel Karimova, Antoine Gedeon, Hélène Munier-Lehmann

**Author notes:** To whom correspondence should be addressed: Hélène Munier-Lehmann. Structural Motility, UMR 144 CNRS/Curie Institute, PSL Research University, Paris, France.

## Abstract

Purine nucleotide biosynthesis is a crucial metabolic pathway responsible that produces building blocks essential for a plethora of cellular processes. In bacteria, the *de novo* purine nucleotide biosynthetic pathway (DNPNB) involves fifteen chemical steps catalysed by fourteen different enzymes. While the mammalian orthologues have been extensively shown to interact and form a metabolon named “purinosome”, the possible existence of a prokaryotic equivalent was only recently revealed for the case of *Escherichia coli*. In this study, we explored the potential conservation of a bacterial purinosome-like complex in *Pseudomonas aeruginosa*, an opportunistic pathogen known for its high antibiotic resistance. Using a bacterial two-hybrid system, we mapped protein-protein interactions among all tested DNPNB enzymes in *P. aeruginosa* and revealed a dense interaction network. An *in-silico* protein-protein docking approach on three core enzymes allowed the structural reconstitution of a complex composed of PurK, PurE and PurC with a 4:8:8 stoichiometry, respectively. Interestingly, a tunnel connecting the different active sites has been revealed, showing a metabolon-like property for possible efficient substrate channelling. These findings support a conserved regulatory organization of purine biosynthesis in bacteria, providing deeper insights into bacterial metabolism and paving the way for potential antibiotic targets.

## 1. Introduction

Purine nucleotide biosynthesis is a fundamental metabolic pathway responsible for the production of purine nucleotides, essential for nucleic acids synthesis and involved in various cellular processes [1]. Nucleotide pools are ensured by two distinct routes: the salvage pathway, and the *de novo* biosynthetic pathway. Both share 5’-phosphoribosyl-1-pyrophosphate (PRPP) as a common precursor and are tightly regulated to maintain a balance between adenylate and guanylate nucleotide pools, as well as optimal energy charges throughout the cell cycle [2].

The *de novo* purine nucleotide biosynthetic (DNPNB) pathway comprises a series of enzymatic reactions that convert PRPP into inosine 5′-monophosphate (IMP), a metabolic branching point that can be further transformed into either adenine or guanine nucleotides via two bifurcated routes, each consisting of two distinct steps [3]. While the general organization of the DNPNB pathway is relatively well-conserved in mammals and bacteria (Fig. 1, Supplementary Fig. S1 and Supplementary Table S1), several notable key differences exist [1, 4, 5]. In bacteria, PRPP is converted into IMP through twelve possible reactions catalysed by eleven different enzymes (Supplementary Table S1), including one bifunctional enzyme (PurH, equivalent to ATIC in mammals). In contrast, mammals utilize ten reactions catalysed by six enzymes (Fig. 1 and Supplementary Table S1), including one trifunctional enzyme (TrifGART) and three bifunctional enzymes (PAICS, ATIC, ADSL). Compared to mammals, the third step in bacteria can follow two routes: i) one involves the folate-dependent enzyme PurN, analogous to the GAR Tfase domain of mammalian TrifGART; ii) the second involves the formate-dependent, prokaryote-specific enzyme PurT. Additionally, the sixth step in prokaryotes is divided into two sequential reactions (Supplementary Fig. S1): the first reaction is catalysed by PurK, an enzyme with no equivalent in mammals and the second by PurE, corresponding to the CAIR synthetase domain of PAICS in mammals.

**Fig. 1.**
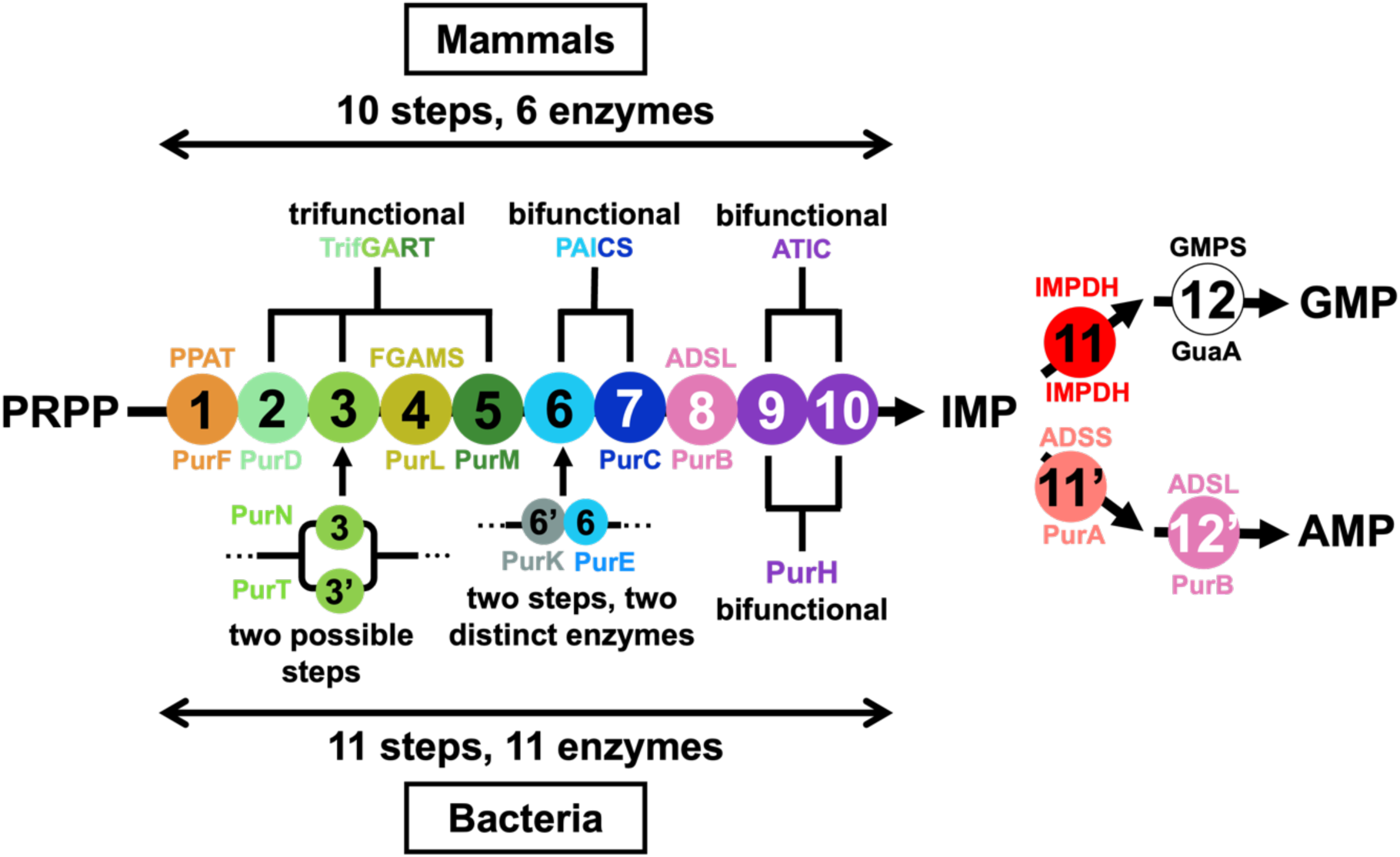
Schematic representation of the DNPNB pathway in mammals and bacteria. The chemical steps involved in the conversion of PRPP into AMP and GMP have been sequentially numbered and given a specific colour. The acronym/name of each enzyme is indicated above the corresponding step number for humans and below the number for bacteria, with the color-coding of the step numbers maintained for the enzyme names.

The purinosome, a metabolon facilitating purine biosynthesis, has been previously discovered in human cells and has gained significant interest. Benkovic’s research group originally identified a colocalization of the first six DNPNB enzymes under purine-depleted conditions in HeLa cells with fluorescence microscopy [6]. Since then, all DNPNB enzymes have been shown to interact within the purinosome [1, 7]. Among the first six enzymes of the DNPNB pathway, the Tango reporter system used for screening protein-protein interactions in cells has revealed that the composition of the purinosome central core comprises enzymes PPAT, FGAMS, and TrifGART, which individually interact with PAICS, ATIC, and ADSL (Supplementary Fig. S1 and Supplementary Table S1) [8]. A more recent study has suggested a revised organization of the purinosome, placing PAICS at the core of the complex [9]. In this same work, they have further showed that the disruption of protein-protein interactions among the purinosome negatively impacts DNPNB intermediate and product pools, confirming the importance of these interactions for its proper functioning of this metabolic pathway [9]. Furthermore, mitochondria-proximal purinosome, termed “active DNPNB metabolon”, has been shown to enhance the efficiency and regulation of this crucial pathway by bringing together all the DNPNB enzymes (and not only the first six ones) in a spatially organized manner [10–12]. The purinosome formation is thus regulated in space and time through various mechanisms, demonstrating that it has a multitude of partners forming a complex functional network rather than being an isolated entity in the cell [7, 13].

Purinosomes have been well-characterized in human cells, reversely less is known with bacteria. Gedeon *et al.* firstly reported in 2023 the existence of a dense interaction network between all DNPNB enzymes in *E. coli* [14]. Using the bacterial adenylate cyclase two-hybrid (BACTH) approach, pairwise interactions between all *E. coli* DNPNB enzymes of this pathway led to the establishment of an intricate network, with PurK, PurE, and PurC as central interactants. Disruption of interactions between PurK and some of the other enzymes has shown to be causative of nucleotide concentrations imbalance and bacterial fitness perturbation, supporting the idea that *E. coli* may also regulate purine biosynthesis through a functional supramolecular assembly. However, the existence and characterization of such organization in other prokaryotes, particularly in pathogenic bacteria, remains to be studied. Given the structural and functional conservation observed in other metabolic complexes among all domains of life, it is reasonable to propose that similar complexes may also be present in prokaryotes.

In this study, we aimed to extend the understanding of purine biosynthesis in a pathogenic bacterium to highlight potential new targets for antibiotic development. Specifically, we have addressed the conservation of this complex in *Pseudomonas aeruginosa*, a notable opportunistic pathogen known for its metabolic versatility and resistance to antibiotics [15]. We followed the same strategy as the one used for the *E. coli* enzymes [14], employing the BACTH approach. Our work revealed extensive binary interactions that link most of the enzymes involved in the DNPNB pathway of *P. aeruginosa* [14]. Furthermore, this study aimed to get insight into the organization of the complex at the molecular level using *in silico* approaches to predict the potential assembly of the core enzymes PurK, PurE and PurC, leveraging the conserved nature of bacterial DNPNB enzymes.

## 2. Material and methods

### 2.1. Bacterial strains, plasmids and composition of culture media

*E. coli* DHT1 strain [16] was used for cloning experiments. The *E. coli* K-12 mutant strains deleted of the genes of interest coding for the DNPNB enzymes obtained from the Keio collection [17] were used for bacterial complementation assays. *E. coli cyaA* strain BTH101 [18] was used for BACTH complementation assays.

Plasmids used in this work are listed in Supplementary Tables S2 and S3 and were assembled by the Gateway^®^ cloning method (Thermo Fisher Scientific and Invitrogen). The BACTH constructs of the vectors harbouring *ftsA* from *E. coli* and *purK* from *P. aeruginosa* were described previously [14, 18].

Open reading frames (ORFs) of all other genes of interest were obtained as synthetic DNA fragments from Twist Biosciences (Sans Francisco, CQ, USA) inserted into the Gateway^®^ entry vector pTwist-ENTR and flanked by attB sequences. Each ORF was subsequently transferred to the attR flanked destination vector pDST25-DEST or pUT18C-DEST using the LR clonase II enzyme mix (Gateway® LR Clonase™ Enzyme mix, Invitrogen). This process enabled the in-frame fusion of ORFs with the sequences encoding T25 or T18 fragments of the catalytic domain of *Bordetella pertussis* adenylate cyclase [19]. All recombinant plasmids have been verified by DNA sequencing (Eurofins Genomics, Les Ulis, France).

The media used in this study were lysogeny broth (LB) and the minimal medium M63B1, whose compositions are as follows: 10 g.L^−1^ peptone, 5 g.L^−1^ yeast extract, 10 g.L^−1^ NaCl, pH 7 and 13.6 g.L^−1^ potassium hydrogen phosphate, 0.2% ammonium sulfate, 0.01% magnesium sulfate, 0.025‰ iron sulfate, 0.002‰ thiamine hydrochloride, 0.2% glucose, pH 7, respectively. For plates, 15 g.L^−1^ of agar was added to the medium.

### 2.2. Bacterial complementation assay

Complementation assays were carried out using the *E. coli* K-12 *pur* and *guaB* mutant strains from the Keio collection [17]. Each mutant deleted of one ORF of interest was transformed with the plasmids pDST25 or pUT18C containing the corresponding deleted ORF from *P. aeruginosa*. The resulting transformants were spread on LB-agar plates (as growth control) or on M63B1-agar plates with the selection antibiotics (kanamycin, and spectinomycin for pDST25 or ampicillin for pUT18C). Each mutant was also transformed with pDST25-*icd* or pUT18C-*icd* as negative controls to demonstrate complementation selectivity. Plates were incubated for bacterial growth overnight at 37°C.

### 2.3. BACTH assay

The BACTH assay for evaluation of protein-protein interactions was carried out following the protocol as described previously [20]. Briefly, enzymes were fused to the T25 or T18 fragments of *B. pertussis* adenylate cyclase and co-expressed in the *E. coli* βTH101 chemical competent strains. Co-transformations were performed with corresponding pST25 and pUT18C vectors (see Supplementary Table S3), and the transformed bacteria were plated on LB-agar supplemented with selection antibiotics, 0.5 mM isopropyl β-D-1-thiogalactopyranoside (IPTG), and 40 µg/mL 5-bromo-4-chloro-3-indolyl-β-D-galactopyranoside (X-gal). Plates were then incubated at 30°C for 36 to 72 hours. Eight single colonies per recombinant protein-pair were subsequently selected and grown in liquid medium overnight in 96 Deep Well plates. After a five-fold dilution, 200 µL was used to measure the optical density at 595 nm (OD_595_) and 200 µL was used to perform β-galactosidase assay by recording ortho-nitrophenol formation at 405 nm (OD_405_). The β-galactosidase activity was measured for six minutes on a Sunrise plate reader (TECAN, Männedorf, Switzerland) and was calculated in arbitrary units (AU) according to the following formula: 1000 x (OD_405_ - OD_405_ blank) / 6 (OD_595_ - OD_595_ blank). Data analysis was performed using GraphPad Prism 10.0 software (San Diego, CA, USA). Cytoscape 3.7.2 [21] was used for representation of the interaction network. Statistical significance was determined by one-tailed t-tests comparing T18 and T25 fragments activities. *p*-values from these comparisons were combined using Stouffer’s method. To further control the level of false discoveries, *p*-values were subsequently adjusted for multiple comparisons using the Benjamini–Hochberg procedure to get an adjusted *p*-value (a.*p*-value; Supplementary Table S4).

### 2.4. Experimental 3D Structures

The 3D structures of proteins and their binding partners were obtained from the RCSB PDB database. In the benchmarking studies, the experimental 3D structures of binding partner complexes were analyzed using the software package Biovia Discovery Studio version 2024 (Dassault Systèmes, San Diego, CA, USA) to identify the interactions between proteins and generate a list of interacting residues. Each complexed structure was then separated into individual PDB files for the protein and its binding partners. These structures underwent preprocessing, including energy minimization, before molecular docking, performed using ZDOCK [22].

Residues identified during the interaction analysis of the complexed structures were classified as either active or passive residue restraints. Active residues refer to surface residues directly involved in the interaction between the protein and its binding partner. Passive residues, on the other hand, do not participate in the interaction and are typically identified based on orthologous complexes or proteins. In this study, only active residues were used as docking parameters to guide the sampling stage, during which the rotational and translational sampling of one protein (the receptor protein) was carried out relative to another stationary protein, which in this case was its binding partner. Before employing a stepwise reconstitution strategy for model refinement, the initial structural modeling of the PurK-PurE-PurC complex was performed using AlphaFold3 [23] using the *E. coli* sequences to predict potential interaction interfaces.

### 2.5. Protein Structure Modelling

For “unbound” docking, the 3D structures of proteins were modeled using both template-based and non-template-based modeling approaches, as described in Abramson *et al.* [23]. For the interacting receptor binding partners, template-based modeling was performed using the software package Biovia Discovery Studio version 2024 (Dassault Systèmes, San Diego, CA, USA). The best final model was selected for docking.

### 2.6. Protein Structure Processing

All PDB structures were prepared using the “Prepare Protein” option in the software package Biovia Discovery Studio version 2024 (Dassault Systèmes, San Diego, CA, USA) with default parameters. The structures were checked for missing atoms, atom names, alternate conformations, incomplete residues, and bond orders. Missing internal loop regions were reconstructed, hydrogen atoms were added, and the final structure was protonated by calculating protein ionization states and residue pKa values. Energy minimization was performed for each structure, with a maximum of 50,000 steps, and rmsd gradient of 0.001 using the Generalized Born implicit solvent model.

For “bound” complexes, the proteins were separated prior to energy minimization. Interacting residues at the interface between effector proteins and receptor binding partner complexes were identified using the software package Biovia Discovery Studio version 2024 (Dassault Systèmes, San Diego, CA, USA) and integrated plugins, with a cutoff distance of 4.0 Å applied to determine interacting residues. For “bound” structures, the ligand’s Euler coordinates were randomized using the software package Biovia Discovery Studio version 2024 (Dassault Systèmes, San Diego, CA, USA) before proceeding with molecular docking.

### 2.7. Protein–Protein Docking

Protein–protein docking was generally performed in two main stages: an initial search and refinement. In the initial search stage, also referred to as sampling, a large number of potential docking (binding) poses were predicted using the docking program. During the subsequent refinement stage, the top-ranked docking poses from the initial search were refined and re-ranked using a more detailed scoring function.

Rigid docking was carried out using ZDOCK [22]. ZDOCK is a rigid docking program that employs 3D grid-based spatial searches during the sampling of binding poses, utilizing an efficient fast Fourier transform (FFT) method [22]. In rigid body docking, the bond angles, bond lengths, and torsion angles of both interacting proteins are kept fixed throughout the process. ZDOCK performs an exhaustive rigid-body search within the six-dimensional rotational and translational space, generating many predicted docking poses.

Each docking pose was assigned a ZDOCK score, comprising pairwise shape complementarity (PSC), desolvation, and electrostatic energy terms. PSC evaluates the total number of atom pairs within a specified distance cutoff between the two proteins, with penalties applied for steric clashes involving core–core, surface–core, and surface–surface overlaps [22, 24]. Desolvation scoring is based on atomic contact energy (ACE), which measures the free energy change associated with breaking protein atom–water contacts and replacing them with protein atom–protein atom and water–water contacts. The total desolvation score was computed as the sum of ACE scores for all receptor–ligand atom pairs within a cutoff distance of 6.0 Å. Electrostatic energy was calculated using Coulomb’s law, incorporating the receptor’s electrostatic potential and the partial charges of ligand atoms.

The docking pose with the highest ZDOCK score was deemed to have the optimal combination of PSC, desolvation, and electrostatic energies.

ZDOCK simulations were conducted using the software package Biovia Discovery Studio version 2024 (Dassault Systèmes, San Diego, CA, USA). Default settings included an angular step size of 6 degrees, generating 26,000 docking poses. Each ZRANK scoring component was reported, and advanced settings for filtering initial sampling poses based on electrostatics and desolvation energies were enabled. No blocked residues or residues known not to form interactions (“passive residues”) were included for either the receptor protein or its receptor binding partners.

Docking poses were filtered using binding site residues or other residues known to form interactions (“active residues”) in both the receptor and ligand partner [25]. The receptor’s binding interface was defined as the set of receptor residues with atoms within 10.0 Å of at least one ligand atom. Similarly, the ligand’s binding interface was defined as the set of ligand residues with atoms within 10.0 Å of at least one receptor atom. Only heavy atoms were considered during this analysis. A docking pose needed to include all active residues from both the receptor and ligand to pass the filtering criteria.

The top 2,000 poses ranked by ZDOCK score were subsequently re-ranked using ZRANK, which incorporates more detailed scoring components, including electrostatics, van der Waals, and desolvation energy terms [26].

For benchmarking “unbound” complexes, simulations were divided into three categories based on the types of unbound structures used: i) experimentally determined 3D structures for both receptor and binding partner: “monomer” structures were derived from protein complexes and docked against receptors or binding partners originating from a different orthologous complex (e.g., *E. coli* or human), both structures were determined in their monomeric forms; ii) hybrid docking: either the receptor or binding partner was modeled, while the other had an experimentally determined 3D structure; iii) fully modeled docking: both the receptor and its binding partner were represented as predicted models.

### 2.8. Binding channel determination

Binding channels were determined using the software package Biovia Discovery Studio version 2024 (Dassault Systèmes, San Diego, CA, USA) plugin based on cavity search algorithm and various cavity were linked according to ligand/protein binding residues determined on PDB orthologs structures: PDB code 7ALE for the human PAICS; PDB codes 1D7A and 2GQS and for the *E. coli* PurE and PurC, respectively).

## 3. Results

A genetic approach based on the BACTH system [27] was employed to explore pairwise interactions among the DNPNB enzymes of *P. aeruginosa*.

The ORFs encoding the enzymes involved in the DNPNB pathway from *P. aeruginosa* were cloned using the Gateway® cloning-adapted BACTH system [19], facilitating their direct insertion in frame with either the T25 (pST25 plasmid) or the T18 (pUT18C plasmid) fragment of the CyaA toxin from *B. pertussis* (see Supplementary Table S3). First, we have checked the expression and the functionality of the fusion proteins. We have taken advantage of the essentiality of all the genes coding for the DNPNB enzymes for bacterial growth in minimal medium (except those encoding PurN and PurT due to the redundancy of these enzymes and PurB as it is encoded by an essential gene). Thus, functional complementation assays were performed using *E. coli* mutants deleted for each gene of interest and sourced from the Keio collection [17]. Mutants were transformed with the corresponding plasmid encoding the relevant T25 or T18 fusions (see Supplementary Table S3). Growth on LB-agar (as a growth control) and on minimal M63B1-agar media was evaluated. As shown in Fig. 2, results show that the expression of the proteins fused either to T25 and to T18 fragments successfully restored the growth of the corresponding *E. coli* mutants on M63B1 minimal medium, meaning that all fusion heterologous proteins are catalytically functional in *E. coli*.

**Fig. 2.**
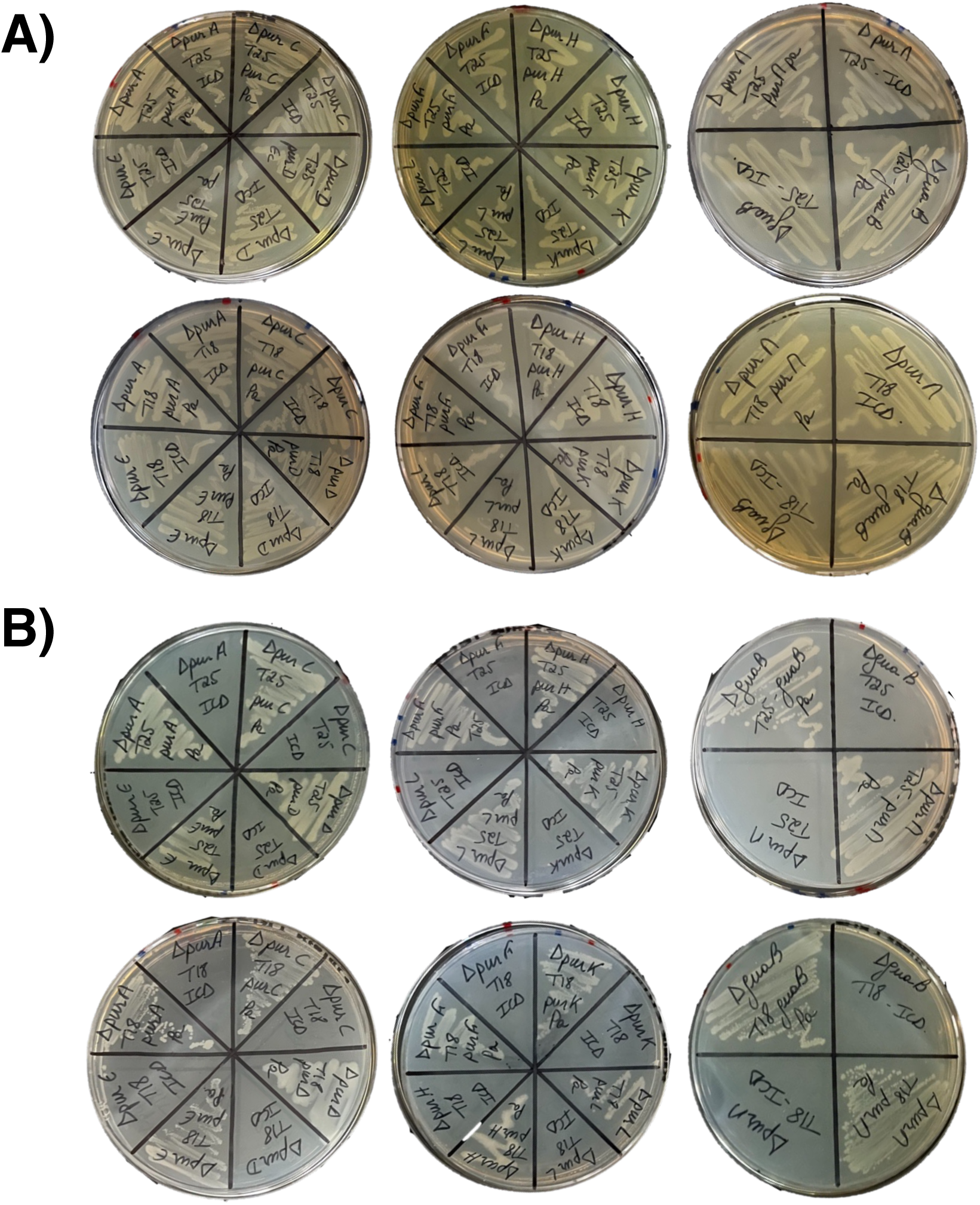
Functional analysis of the T25- and T18-fused proteins. Each mutant strain from the Keio collection [17] was transformed with either pDST25 or pUT18C plasmid derivatives containing the corresponding missing gene. The transformants were then plated on LB-agar (growth control, panel A) and M63B1-agar (panel B). To ensure complementation specificity, the mutants were also transformed with pDST25-*icd* or pUT18C-*icd* plasmids harbouring *icd* gene as negative controls. Due to the salvage pathway, all the tested mutants exhibited growth on LB agar whatever the transformed plasmid (panel A). On minimal media, however, no growth was observed after bacteria transformation with a plasmid harbouring *icd* gene (panel B).

Then, for BACTH assays, screening of protein-protein interactions among DNPNB enzymes was assessed by co-expressing the hybrid proteins in BTH101 bacteria and measuring β-galactosidase activity in liquid cultures (as described in Materials and Methods). Non-fused T25 and T18 fragments (T25-empty/T18-empty) and T25 and T18 fused to the leucine zipper of GCN4 (T25-Zip/T18-Zip) have been used as negative and positive controls, respectively. FtsA (filamentous temperature sensitive A), an *E. coli* cytosolic protein unrelated to the DNPNB pathway, has also been employed as a negative control as described in Gedeon *et al.* [14]. Two partners were considered as interacting when the measured β-galactosidase activity was five-times higher than that of the negative controls (i.e. > 120 AU).

The oligomeric state of all the proteins was verified by examining their homo-interactions (Supplementary Fig. S2). To our knowledge, except for IMPDH [28, 29], no other DNPNB enzyme from *P. aeruginosa* has been purified and characterized. Therefore, due to the lack of information regarding the oligomeric state of these enzymes, data from their *E. coli* homologs, previously described in the literature using classical biochemical approaches and with which they share more than 60% sequence identity (except for PurK, with 38% sequence identity), were used. Our data did show β-galactosidase activity levels (Supplementary Fig. S2) that were consistent with the oligomeric states of the *E. coli* proteins (Supplementary Fig. S1).

Next, a detailed comprehensive screening of all binary interactions between *P. aeruginosa* enzymes was conducted to define an interaction network. Examples of the BACTH system results are displayed in Fig. 3 with the three following baits (as T18- and T25-fusions): PurK, the enzyme with no eukaryotic equivalent (Fig. 3A), PurE (Fig. 3B) and PurC (Fig. 3C), all involved in the catalysis of three successive steps of the DNPNB pathway (steps 6’, 6 and 7, respectively, refer to Supplementary Fig. S1). The rest of the data for all the other enzymes are presented in Supplementary Fig. S3. The ensemble of these results clearly shows the existence of several interaction nodes. For a finer statistical control of false positives, the whole β-galactosidase activity values were further analysed using a two-dimensional statistical test to obtain adjusted merged *p*-values with the Stouffer method coupled with the Benjamini-Hochberg procedure (Supplementary Table S4). These analyses, presented as a two-dimensional matrix (Fig. 4A) and a network map (Fig. 4B), revealed a dense network of significant binary interactions linking all tested enzymes. Indeed, a core was identified, with notably PurE and PurK being hubs of the pathway. PurH, the only bifunctional enzyme of the pathway, is also identified as an important interacting partner. Interestingly, PurK, PurE, and PurC are enzymes that previously showed to have a key central position within the DNPNB megacomplex in *E. coli*. In mammalian cells, PurK is absent, and PurE and PurC are replaced by the bifunctional enzyme PAICS. Because of these interesting evolutionary divergences, we opted to model the structure of the ternary complex PurK, PurE and PurC and analyse the positioning of the different active sites as these enzymes sequentially follow one another in the pathway. The 3D-structures of the *E.* coli PurK, PurE and PurC have been solved by X-ray crystallography (PDB code 1B6S [30], PDB code 1D7A [31] and PDB code 2GQS [32], respectively) and they have been used to model the ones from the *P. aeruginosa* counterparts (Fig. 5). We have chosen to construct the complex (Fig. 6) by a step-by-step strategy by incrementally generating the models [33]. The 3D-structure of the human bifunctional PAICS (PDB code 7ALE) has been used to reconstruct the association between PurE and PurC from *E. coli*. This was possible due to the sequence similarity between *E. coli* PurE and the CAIRS domain of PAICS (25% identity), and between *E. coli* PurC and the SAICARS domain (30% identity). The central core of PAICS is composed of eight CAIRS domains. Similarly, PurE is octameric adopting a cuboid shape with several hydrophobic areas capable of binding potential protein partners on various faces (Fig. 6C). Then, PurC was initially isolated as a dimer, and we proceeded to dock a dimer of PurC onto the octamer of PurE. Similarly to what has been observed in PAICS (PDB code 7ALE) where two SAICARS domains are located on each face of the central core, a dimer of PurC docks onto each of the lateral faces of the cuboid, resulting in four dimers of PurC. However, the study of the binding interface between PurC/PurE shows that only one of the PurC monomers binds correctly to PurE. We therefore questioned the possibility of binding each PurC monomer independently. A study of the binding energies shows better stability of the complex after the independent docking of two PurC monomers compared to a PurC dimer (Fig. 6E). After docking a total of eight PurC monomers onto the PurE octamer, we sought the potential binding site between the PurE/PurC complex and PurK (Fig. 6B). The docking of PurK onto the complex revealed a significant interaction cluster on the faces of the cuboid. Similarly, as before, we docked each PurK monomer onto the PurC/PurE complex and observed the possibility of binding two PurK monomers to each free PurE interface (Fig. 6A-B). Once the complex with a 4:8:8 stoichiometry (PurK/PurE/PurC) was generated, an equilibration was performed to obtain a stable complex. The same organization could be modelled for the *P. aeruginosa* PurK/PurE/PurC complex (Supplementary Fig. S5). One method of validating our complex was the consistency between the generated model and the complex’s ability to perform the conversion reaction of AIR to SAICAIR (Fig. 7A). Interestingly, we observed that the binding sites between PurK and PurE were contiguous, forming a channel that allows the NCAIR formed by PurK to be converted into CAIR by PurE (Fig. 7B). This continuum extends between PurE and PurC, allowing CAIR to in turn be converted into SAICAIR, thus completing the expected reaction from AIR to SAICAIR (Fig. 7). Thanks to this channel and the presence of multiple monomers of the same enzymes, the simultaneous handling of four AIR molecules is conceivable, allowing parallel catalysis of different substrates.

**Fig. 3.**
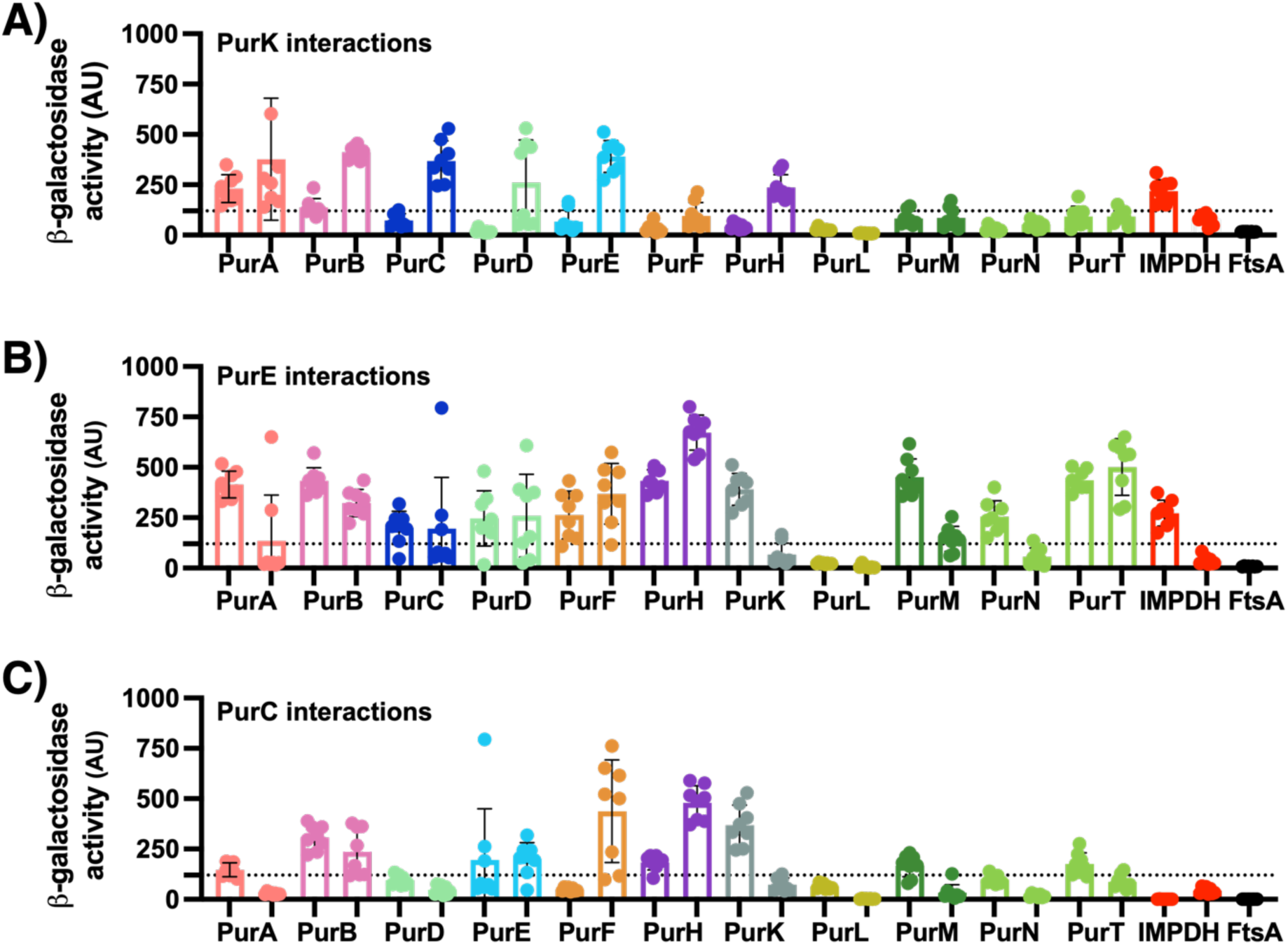
Investigation of the interactions between PurK (panel A), PurE (panel B) and PurC (panel C) and the other DNPNB enzymes using the BACTH system. β-galactosidase activity, expressed in arbitrary units (AU), was measured as detailed in the Materials and Methods section. Each enzyme (identified below the histogram, with colours matching Fig. 1) was tested as a prey in T25 (left) or T18 (right) fusion, with PurK, PurE or PurC acting as the bait (using the corresponding T18 or T25 fusion). FtsA was included as another negative control under the same conditions. The dotted line marks five times the β-galactosidase activity value (120 AU) of the negative controls. Data are presented as means ± SD from eight independent measurements (n = 8).

**Fig. 4.**
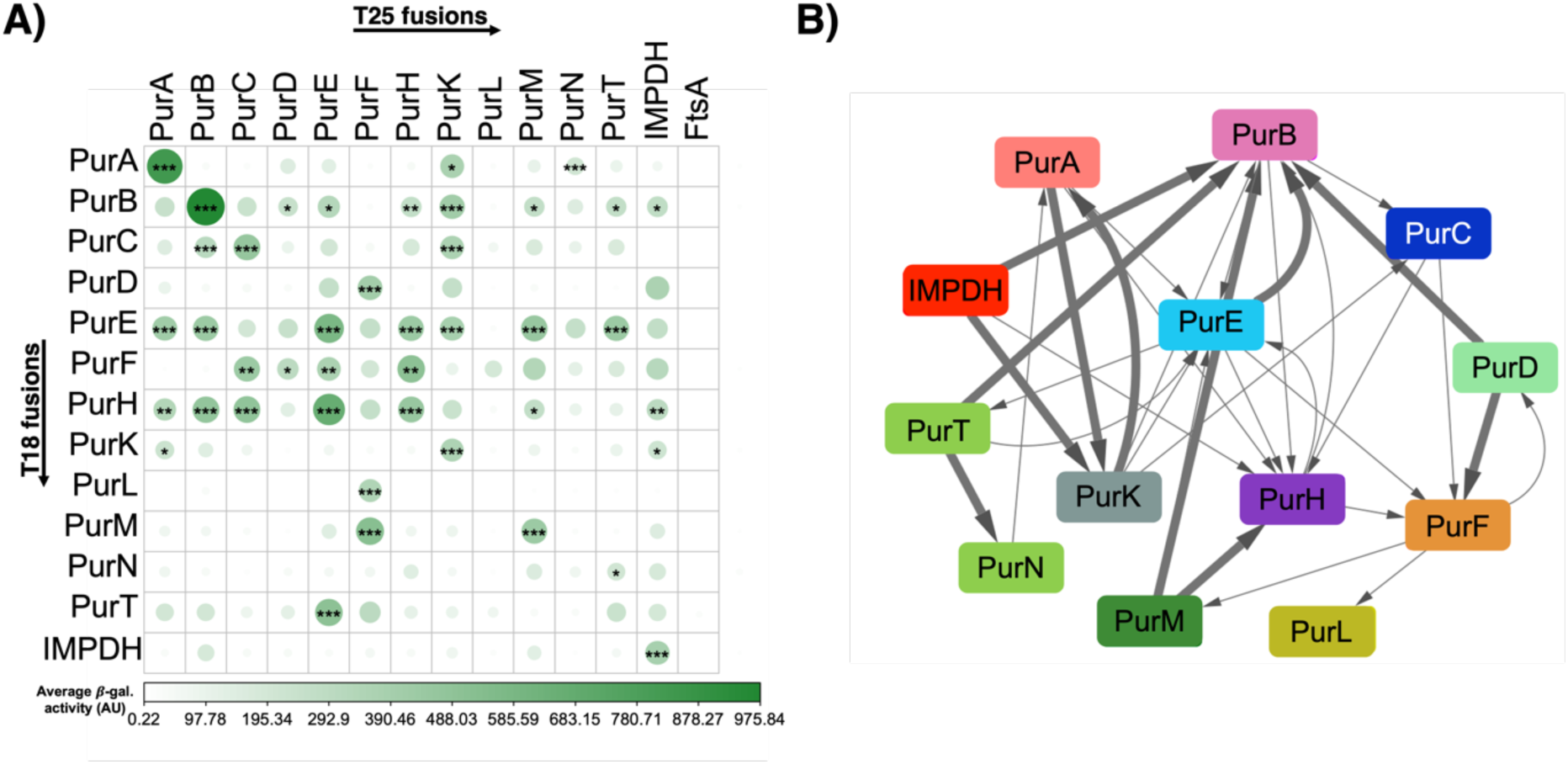
Statistical analysis of the β-galactosidase activity values from BACTH screening. A) Two-dimensional matrix showing the significance of the interactions detected by the BACTH system between the T18 (columns) and T25 (rows) versions of the DNPNB enzymes. The intensity of the green colour and the size of the circles correspond to the β-galactosidase activity value (means from eight independent measurements (n = 8), see Fig. 3 and Supplementary Fig. S3). An adaptive Benjamini–Hochberg correction was applied to obtain an adjusted *p-value*s (a.*p-value*s). *, 1% < a.*p-value* < 5%; **, 0.1% < a.*p-value* < 1%; ***, a.*p-value* < 0.1%. a.p-values are given in Supplementary Table S4. B) Interaction network generated using Cytoscape 3.7 [21] showing enzyme interactions identified by the BACTH system, with each arrow representing a node between T18 and T25 versions. The width of the arrows reflects the corresponding a.*p-value*s. Interactions of each protein with itself (homooligomerization) are not shown in this diagram.

**Fig. 5.**
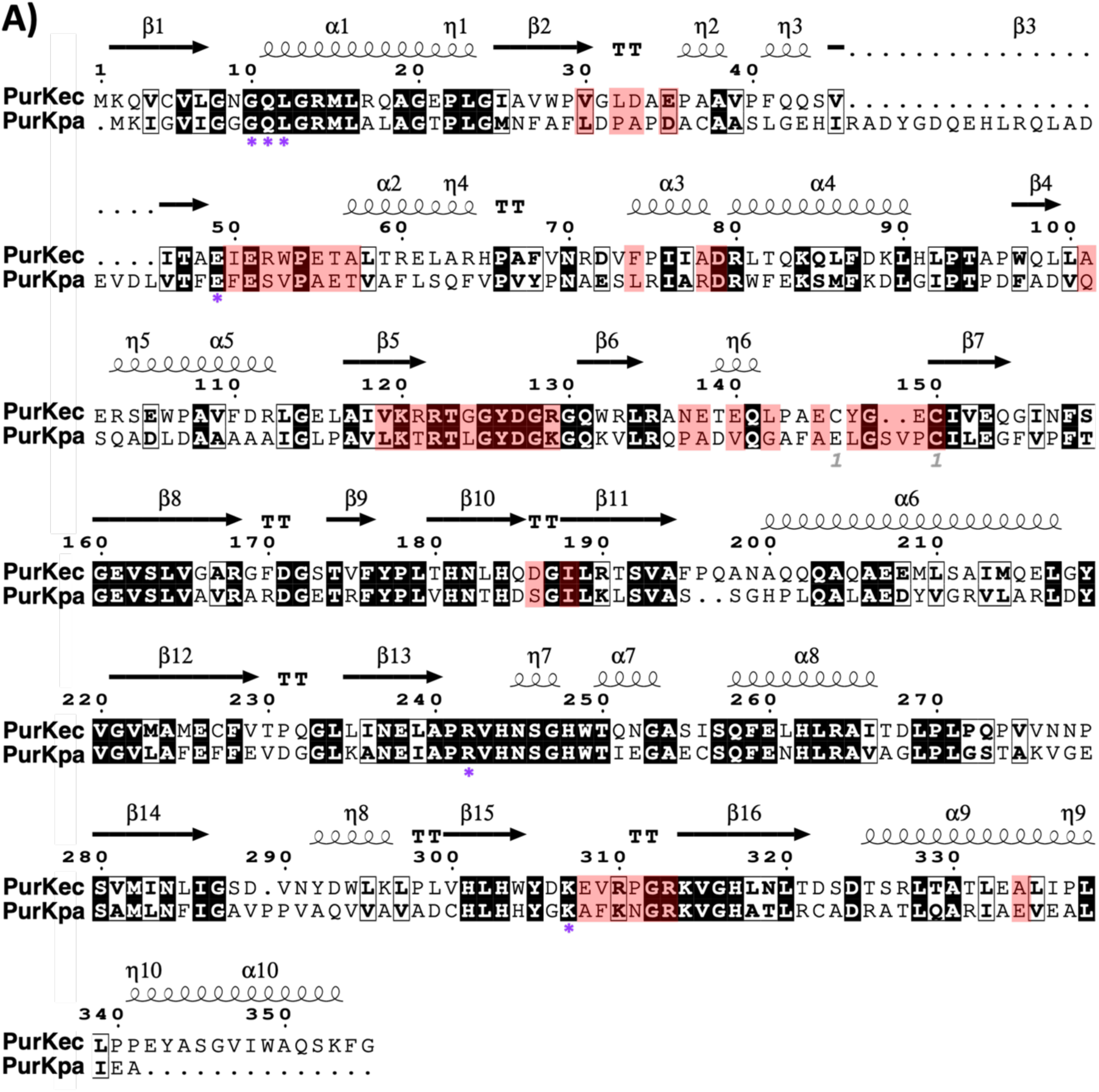
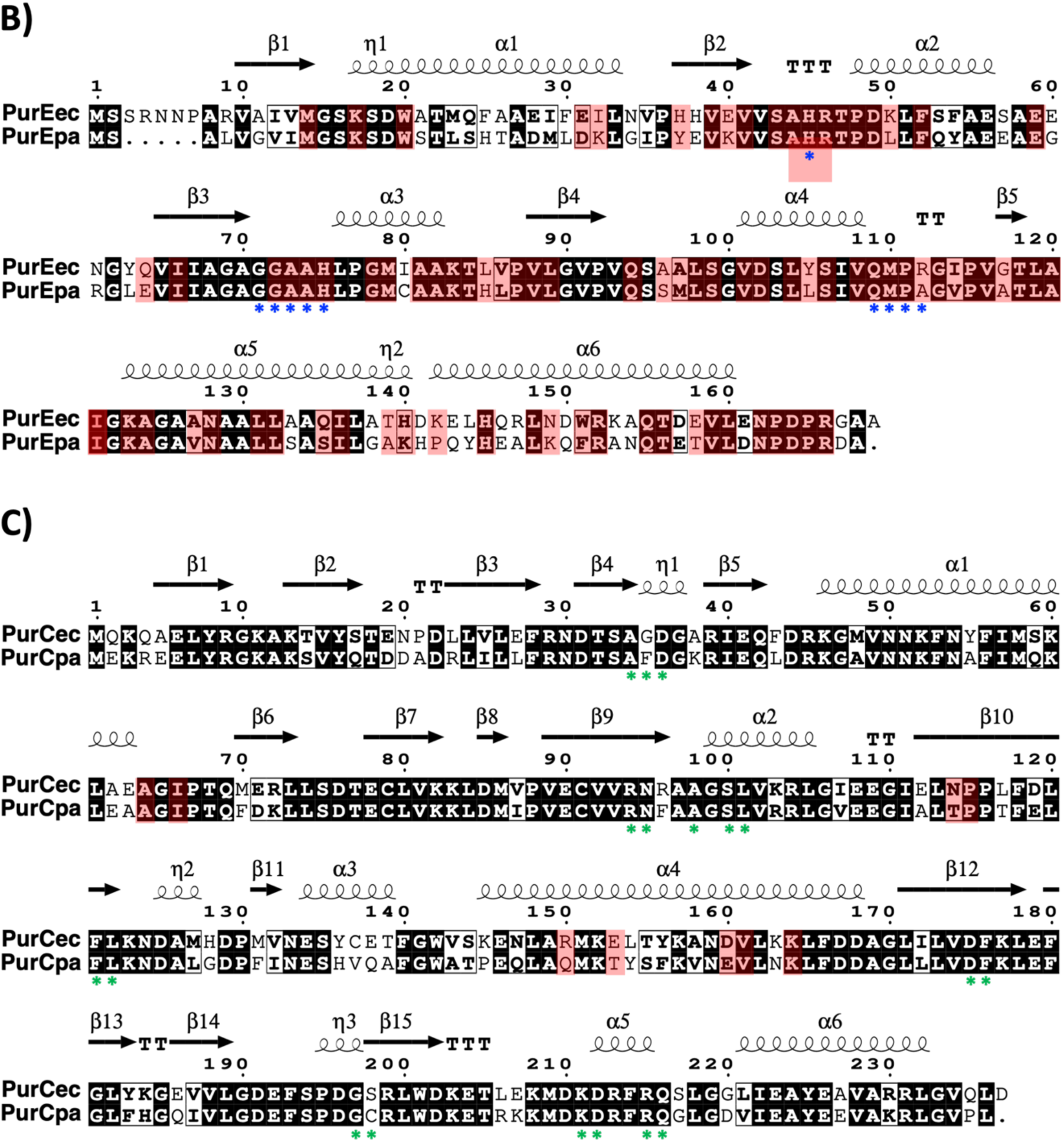
Multiple sequence alignments between *E. coli* and *P. aeruginosa* PurK (panel A), PurE (panel B) and PurC (panel C). Residues highlighted in red indicate interaction surface residues involved in the interactions between PurK, PurC, and PurE. Conserved residues are enclosed in black boxes, while residues involved in the ligand-binding site (catalytic residues) are marked with color-coded asterisks: blue for PurE, green for PurC, and purple for PurK. Sequence alignment was performed using Clustal Omega [51] and visualized with ESPript 3.0 [52].

**Fig. 6.**
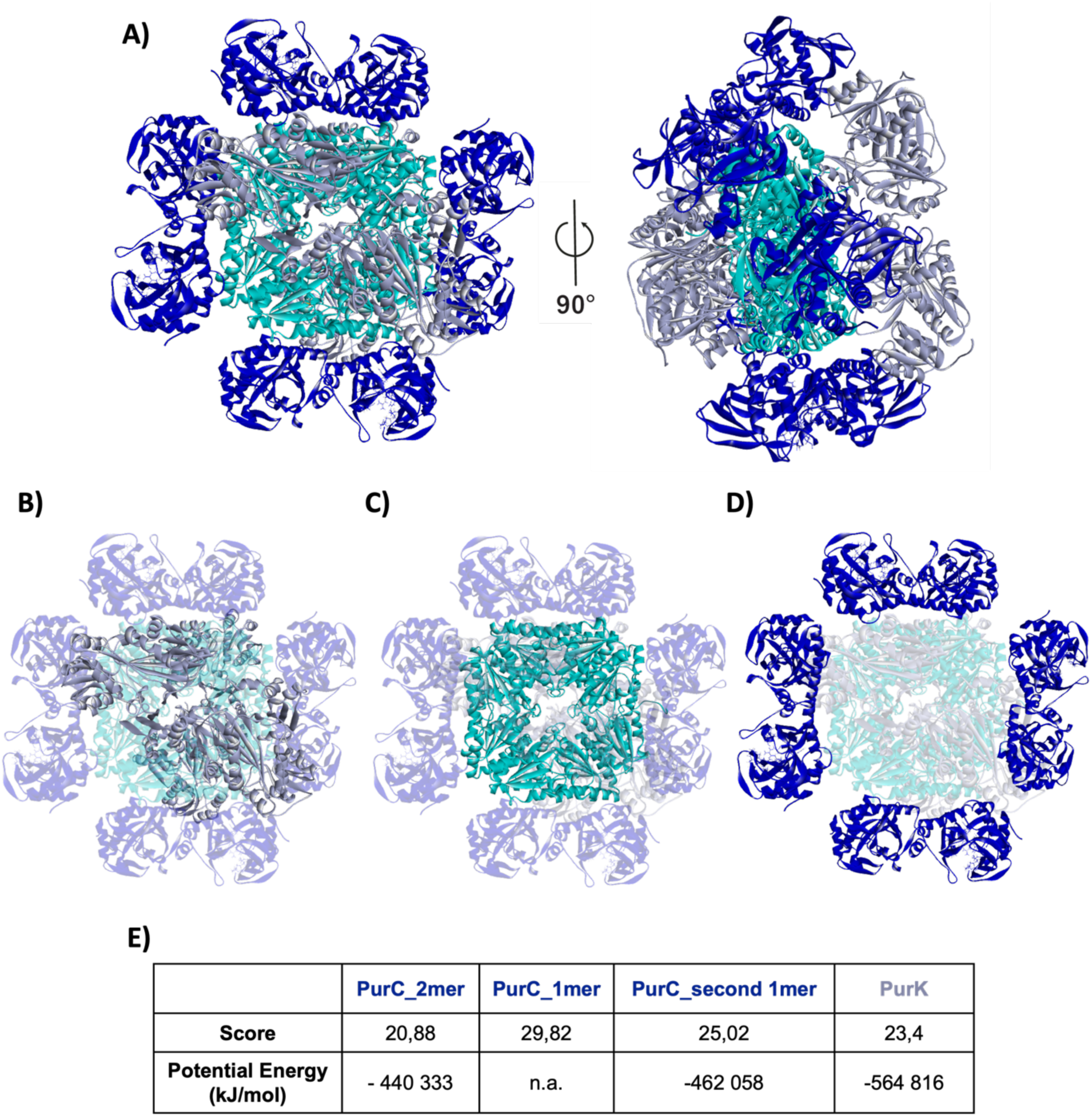
Molecular docking experiments predicting protein-protein interactions between PurK, PurE, and PurC from *E. coli*. A) Predicted complex of the three enzymes, with PurK displayed in grey, PurE in cyan, and PurC in dark blue. B-D) The full complex is shown; B) with PurE and PurC faded, highlighting the PurK dimer positioned on each side of the PurE octamer; C) PurK and PurC faded, emphasizing the octameric structure of PurE; D) PurE and PurK faded, displaying the eight monomers of PurC, with two monomers interacting with each side of PurE. E) Table summarizing docking scores and potential binding energies for interactions involving a PurC dimer (PurC_2mer), a single PurC monomer (PurC_1mer), another PurC monomer in the presence of a bound PurC monomer (PurC_second 1mer), and a PurK dimer (PurK). n.a.: not applicable.

**Fig. 7.**
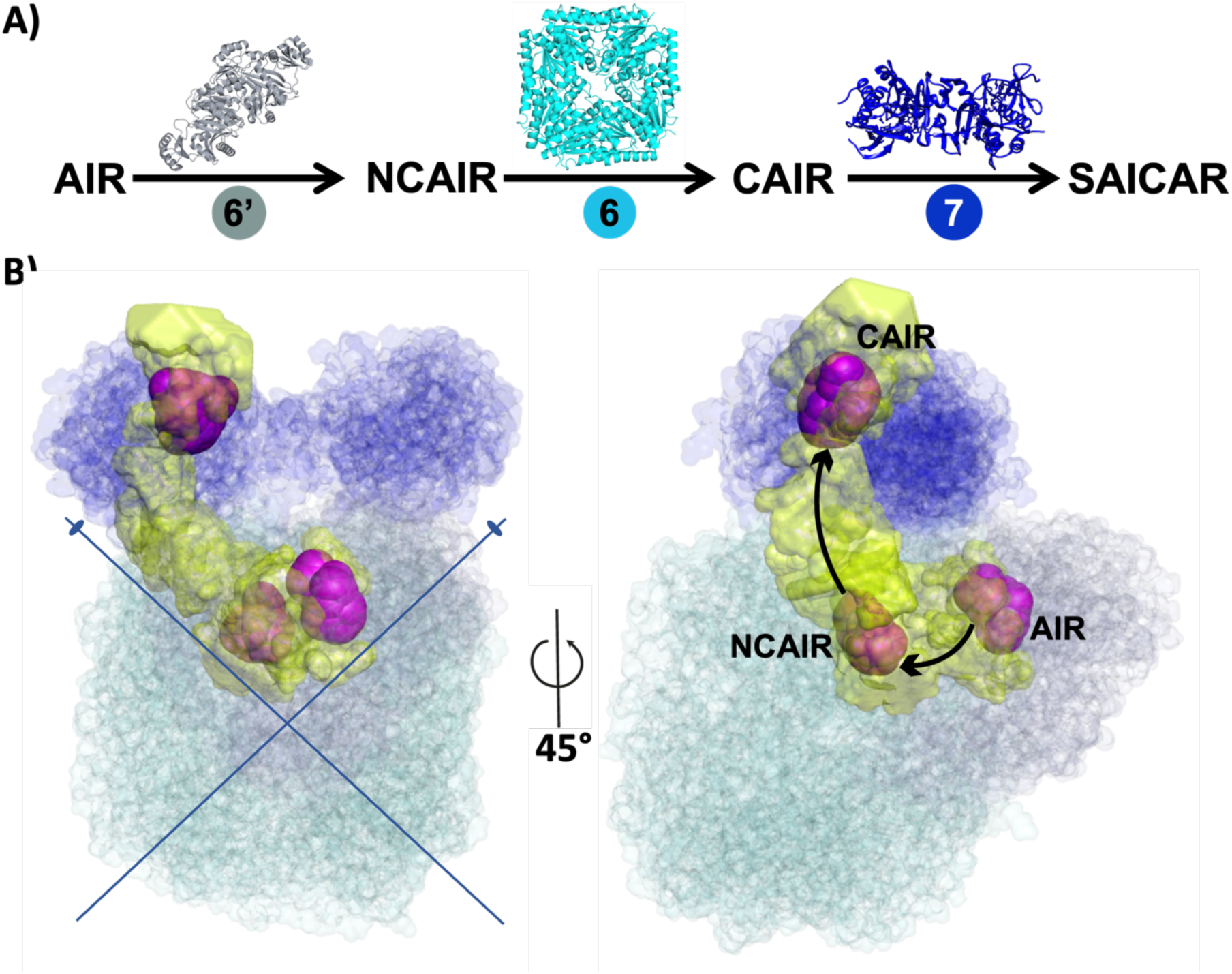
Molecular docking results illustrating substrate channelling between PurK, PurE, and PurC in the *de novo* purine nucleotide biosynthetic pathway. A) Sequential reactions (6’, 6, and 7) catalyzed by PurK, PurE, and PurC, respectively in the DNPNB pathway (Supplementary Fig. S1). B) Predicted channelling pathway, displayed in yellow, between the three enzymes in the previously predicted enzyme complex (see Fig. 6), showcasing their spatial arrangement to support efficient intermediate transfer. Intermediates, displayed in magenta, are transferred across the active sites of the enzymes.

## 4. Discussion

*Pseudomonas aeruginosa* is an opportunistic Gram-negative pathogen, which remains an important public health threat because of the rise of emerging antibiotic-resistant strains [15, 34]. To counteract this phenomenon, it is of utmost importance to discover new potential targets. Previous microbiology work is in favour of purine biosynthesis as a possible promising target in *P. aeruginosa*. For instance, as it is the case for *E. coli* [17], all DNPNB genes (except those coding for PurN and PurT) have been shown to be essential for growth of *P. aeruginosa* in synthetic minimal medium [35]. Moreover, Poulsen *et al.* have selected nine diverse strains of *P. aeruginosa* and assessed their growth in five different growth conditions. Notably, DNPNB genes have proven to be crucial in the three additional growth conditions that closely resemble conditions associated with human infections. A *P. aeruginosa* strain that lost the *purH* gene was not anymore infectious in a neutropenic mouse model. Another research study [36] revealed that a mutant strain, which was deficient in the *purEK* operon and thus required adenine for growth, became avirulent using the same mouse model.

The ensemble of these data supports that DNPNB enzymes could serve as attractive targets for the development of new antibacterial agents aimed at inhibiting their catalytic activity. Alternatively, another promising approach could be to target protein-protein interactions. Indeed, these last years, protein-protein interactions have been of most significant interest for the subsequent development of antibacterial agents targeting specific oligomerisation protein surfaces. Several molecules perturbing fundamental mechanism in pathogenic bacteria, such as cell division or gene expression machineries, have been reported and seem to be promising candidates for the next generation of antibiotics [37, 38].

In this work, the BACTH system [39] has been used to map binary protein-protein interactions of the *P. aeruginosa* DNPNB enzymes. This well-established technique has been employed for the identification and description of several bacterial megacomplexes, such as tricarboxylic acid cycle enzymes from *Bacillus subtilis* [40] or partitioning ParB protein partners from *P. aeruginosa* [41], to name a few. Our present results are in favour of the existence of a dense interaction network between *P. aeruginosa* DNPNB enzymes and constitutes the second evidence of existence of such megacomplex in bacteria [14]. Moreover, the overall interacting network resembles the one we have previously depicted in *E. coli* [14], PurK, PurE and PurH appearing to be involved in most of the interactions. In accordance with our *in vitro* data, a stochastic predictive AI-based analysis of *P. aeruginosa* genome proposed potential interactions between all the DNPNB enzymes (including GuaA), with PurH being the top interactant among these proteins. On the other hand, none of the large-scale analysis have determined interactions among *P. aeruginosa* DNPNB enzymes, probably due to their transient nature. Notably, one proteome-wide analysis *via in vivo* cross-linking coupled to mass spectrometry in *P. aeruginosa* cells did not show any interactions between DNPNB enzymes other than the homo-oligomerization of PurA, PurH and IMPDH [42].

*P. aeruginosa* PurK, PurE and PurC remain orphan proteins that are poorly characterized or have never been studied. Conversely, PAICS, the equivalent of PurE-PurC in human was the object of several studies deciphering its molecular regulation [43] or its implication in genetic defects and cancer [44–46]. However, recent developments in tools and methodologies could open the road for a better understanding of such enzymes [47], especially from bacteria. To get a new further insight into PurK/PurE/PurC interaction, we started with a modelling approach by using AlphaFold3 server [23]. Despite being a powerful tool for protein structure prediction and reconstructions of some protein complexes [48, 49], further developments are needed to gain higher performance and refined accuracy [50]. As such, our attempt to reconstitute the complex regrouping *E. coli* or *P. aeruginosa* PurK/PurE/PurC using AlphaFold3 server proposed a model with a high predicted aligned error (Supplementary Fig. S4) and was incompatible with the solved 3D-structure of the human homolog of PurE/PurC, namely PAICS. This discrepancy arose from several structural inconsistencies in the AlphaFold3 model. When superimposed with the experimentally solved PAICS structure (PDB: 7ALE), the AlphaFold3 model displayed an incorrect arrangement of the ligand channels, suggesting an improper spatial organization for ligand transformation. The symmetry axis was not conserved, leading to a mispositioning of PurC relative to PurE, where only one monomer was engaged in the interaction while the other was significantly displaced. Additionally, interactions in the AlphaFold3 model were predicted only on one face of PurE while potential interactions on the other face were not explored. This further highlights the model’s limitations in accurately capturing the structural organization of the complex. Thus, a step-by-step reconstruction using protein/protein docking approaches was used to generate the bacterial complex. Given the high similarity between *E. coli* and *P. aeruginosa* enzymes, a similar organization is predicted for both bacterial complexes with a 4:8:8 stoichiometry of PurK, PurE and PurC, respectively. Moreover, analysis of the catalytic site arrangements of the three enzymes revealed the presence of a channel that enables ligand transfer across the active sites, thus facilitating efficient and coordinated enzymatic reactions. The identification of a dense interaction network among DNPNB enzymes, combined with structural predictions, strongly supports the hypothesis that these enzymes assemble into a functionally relevant supramolecular complex. Metabolon formation has been demonstrated in humans through different approaches, yet the precise composition and the spatial organization of the purinosome remain to be fully elucidated. Similarly, while our bacterial model provides valuable structural insights, it is currently limited to the core PurK/PurE/PurC complex. Given that our BACTH results identified additional interactions involving other DNPNB enzymes, future studies should aim to expand this model to encompass the full pathway. This broader characterization could reveal novel regulatory mechanisms and identify key protein-protein interaction interfaces within the complex that could serve as potential antibiotic targets. Given the emerging interest in disturbing key protein-protein interactions, further exploration of this complex may uncover new potential drug targets and opportunities to interfere with essential enzymatic assemblies.

## Acknowledgements

We are grateful to Jean-Marc Ghigo for providing the *E. coli* mutant strains from the Keio collection. We thank the “Plateforme de Milieu” of the Institut Pasteur for media culture preparations.

This work was supported in part by the Centre National de la Recherche Scientifique (CNRS), the Institut National de la Santé Et de la Recherche Médicale (INSERM) and the Institut Pasteur. Nour Ayoub acknowledges a PhD fellowship from Médicament, Toxicologie, Chimie et Imagerie Ph.D. school (MTCI, ED 563), Université Paris Cité.

## CRediT authorship contribution statement

N.A., N.P. and H.M.L. conceptualized and designed experiments; H.M.L. supervised the work; N.A. and G. K. performed cloning; N.A. performed BACTH assays; N.P. conducted docking experiments; N.A., N.P., Q.G.G., A.G., and H.M.L. analysed the data; N.A., N.P., A.G. and H.M.L. wrote the manuscript. All authors completed and edited the paper.

## Appendix A. Supplementary data

Supplementary data (Supplementary Figures S1-5, Supplementary Tables S1-4 and references) can be found online.

