## Supplementary data for "Exploring the multi-protein assembly of the enzymes of the *de novo* purine nucleotide biosynthetic pathway from *Pseudomonas aeruginosa*"

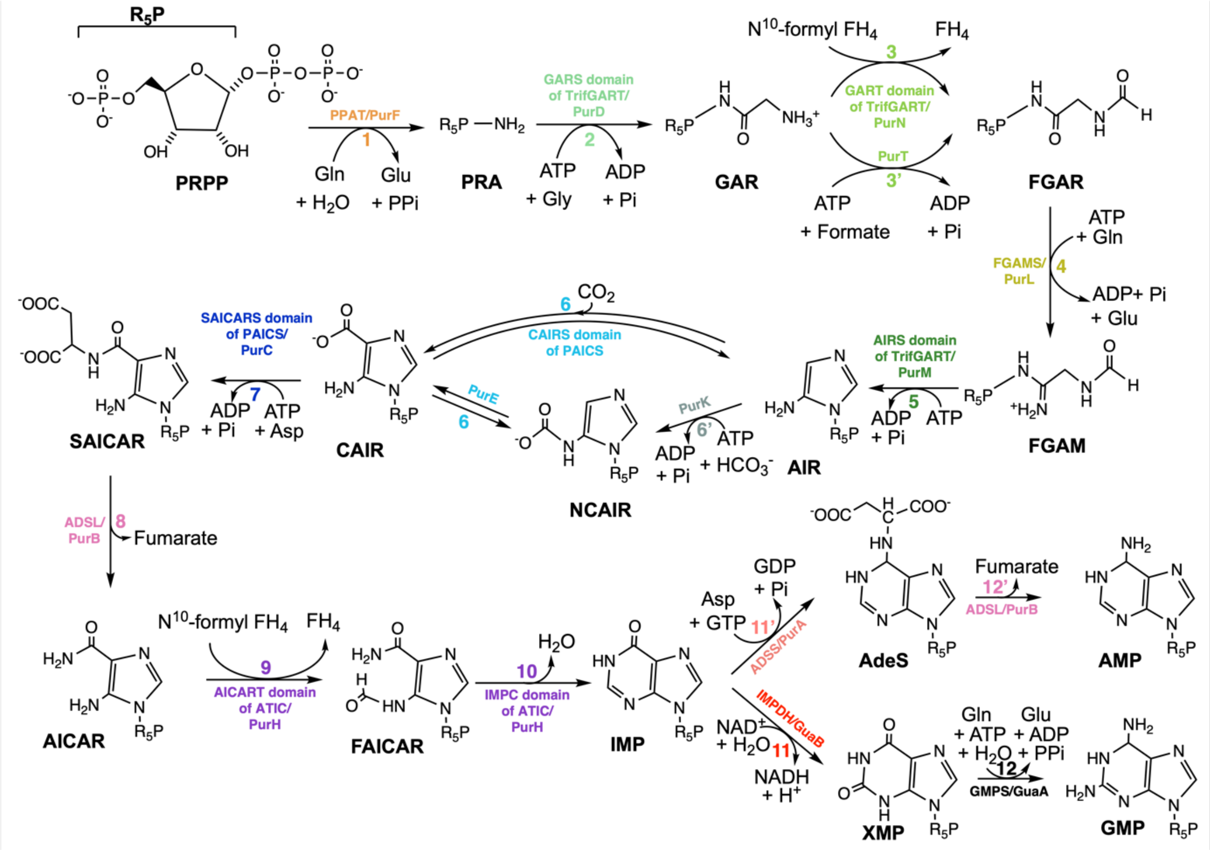


**Figure S1**. *De novo* purine nucleotide biosynthesis (DNPNB) pathway. Representation of the sequential conversion of PRPP into AMP and GMP. The steps are numbered and labeled with specific colors. The acronyms of the enzyme involved in each step is provided for both mammals and *E. coli*, the corresponding full names are listed in Supplementary Table 1.

In vertebrates, six enzymes are responsible for the conversion of PRPP to IMP, which is then transformed into GMP or AMP by two additional reactions. Bacteria require fourteen enzymes to catalyze these different steps.

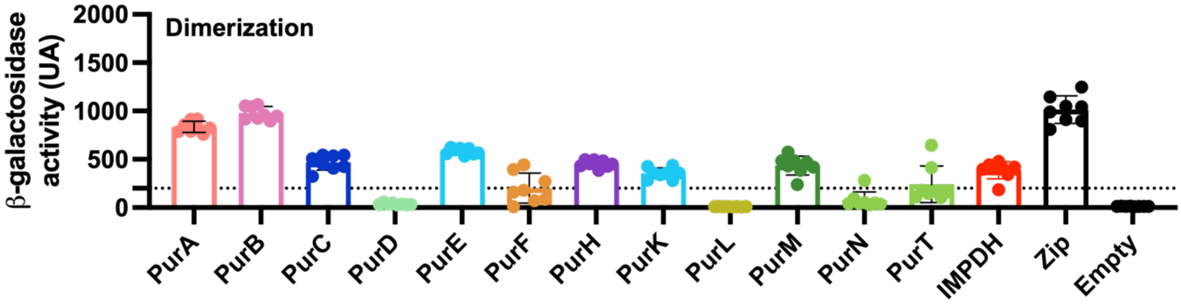


**Figure S2**. Oligomerization status of the different enzymes from the DNPNP pathway from *P. aeruginosa*. Interactions of T25 and T18 fusions of the same enzyme were assayed using the BACTH system. Positive (Zip corresponding to T25-Zip/T18-Zip) and negative (corresponding to T25-empty/T18-empty) controls are also reported. β-galactosidase activities are expressed in arbitrary units (AU).

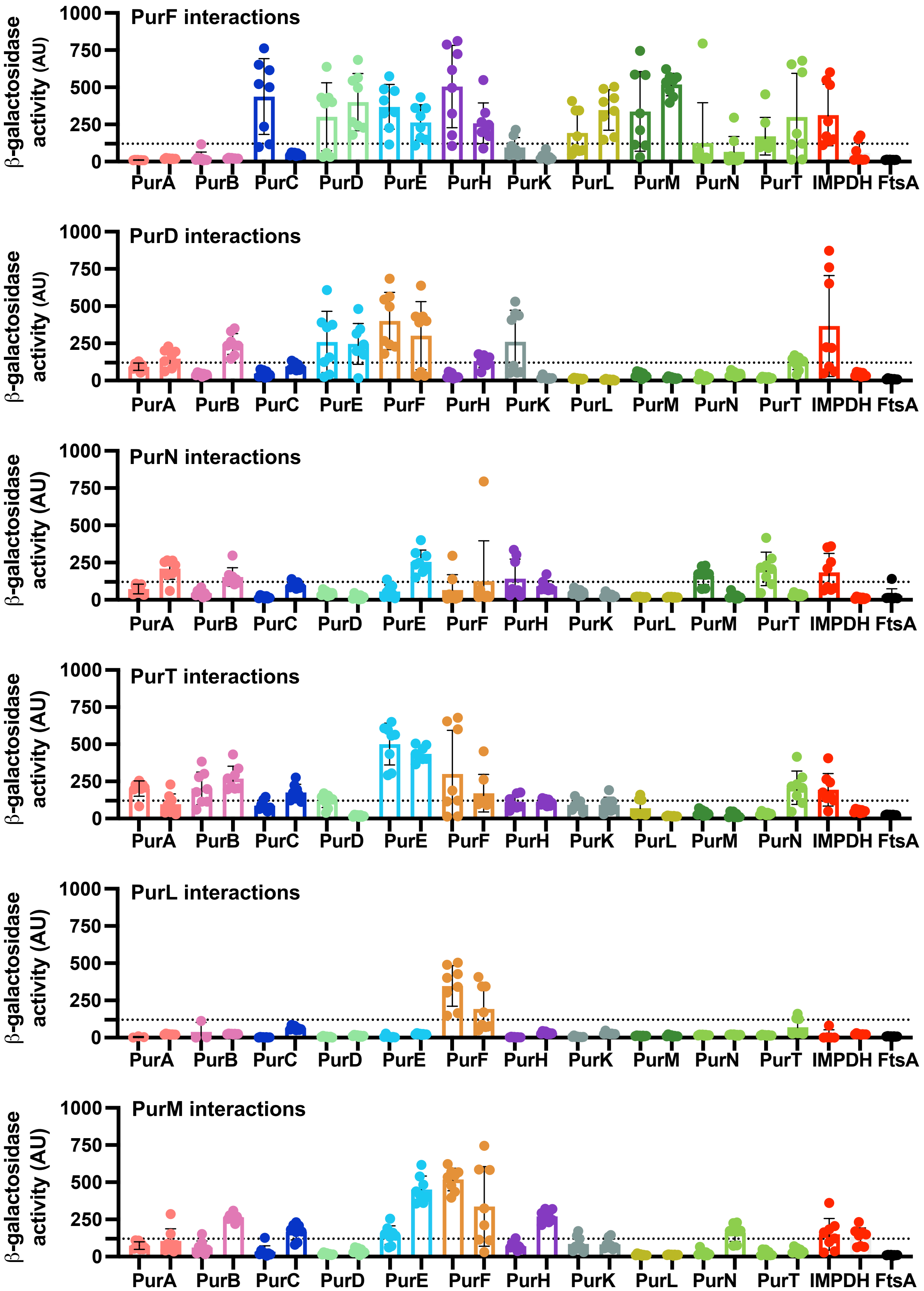

β-galactosidase activity (AU)
1000
750
500
250
0
PurF interactions
PurA PurB PurC PurD PurE PurH PurK PurL PurM PurN PurT IMPDH FtsA
PurD interactions
PurA PurB PurC PurE PurF PurH PurK PurL PurM PurN PurT IMPDH FtsA
PurN interactions
PurA PurB PurC PurD PurE PurF PurH PurK PurL PurM PurT IMPDH FtsA
PurT interactions
PurA PurB PurC PurD PurE PurF PurH PurK PurL PurM PurN IMPDH FtsA
PurL interactions
PurA PurB PurC PurD PurE PurF PurH PurK PurM PurN PurT IMPDH FtsA
PurM interactions
PurA PurB PurC PurD PurE PurF PurH PurK PurL PurN PurT IMPDH FtsA

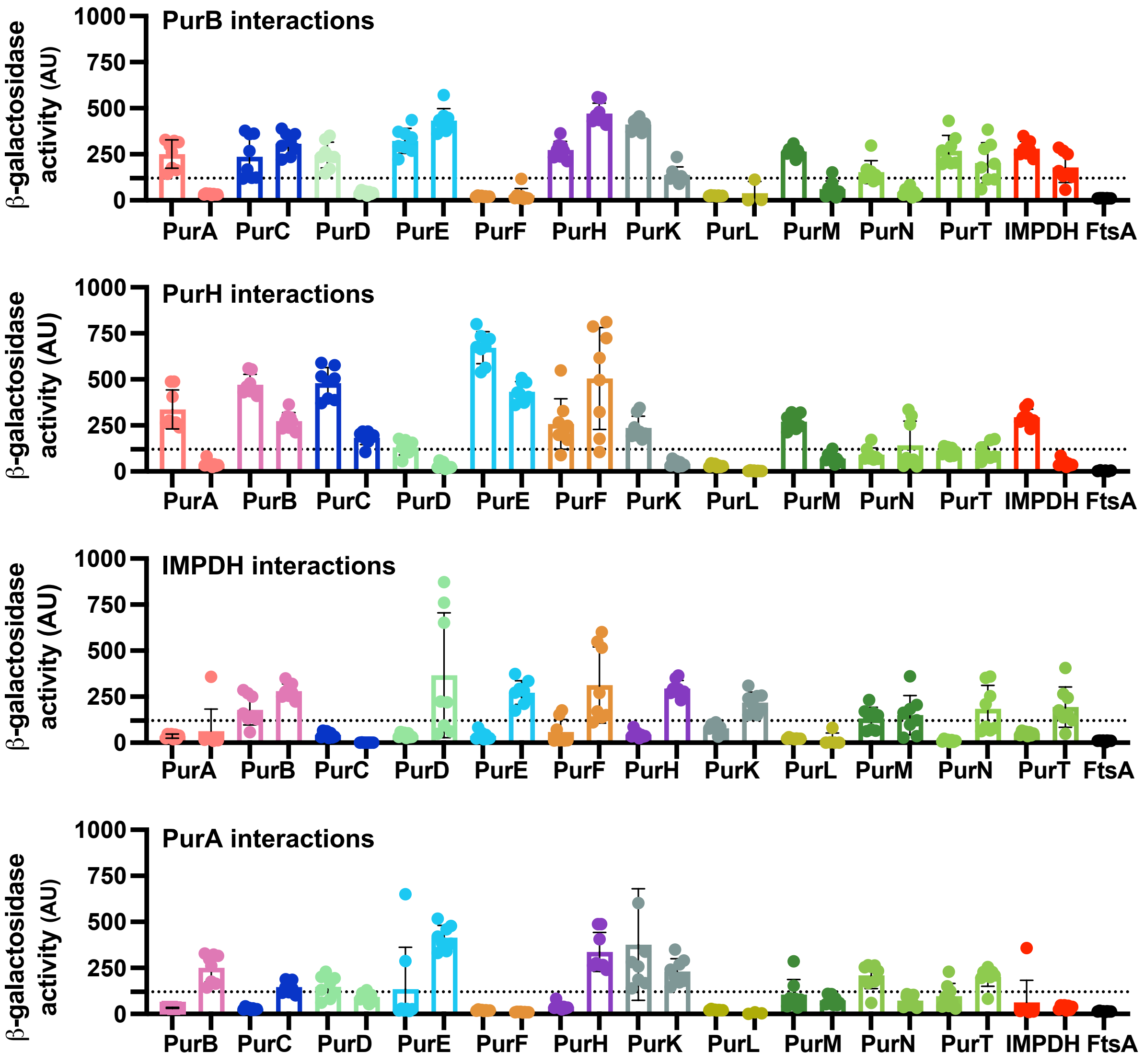


**Figure S3**. Interactions of each *P. aeruginosa* DNPNB enzyme (using T18 or T25 fusion; the name is indicated in bold above each histogram, with the order of the histograms (from top to bottom) corresponding to the step in which the protein is involved, except for PurK, PurE and PurC which data are depicted in Figure 3) taken as a bait and the other proteins of this pathway taken as preys (identified below the histogram) in the T25 (left) or T18 (right) fusion. FtsA was included as another negative control under the same conditions. The dotted line marks five times the β-galactosidase activity value (120 AU) of the negative controls. Data are presented as means ± SD from eight independent measurements (n = 8).

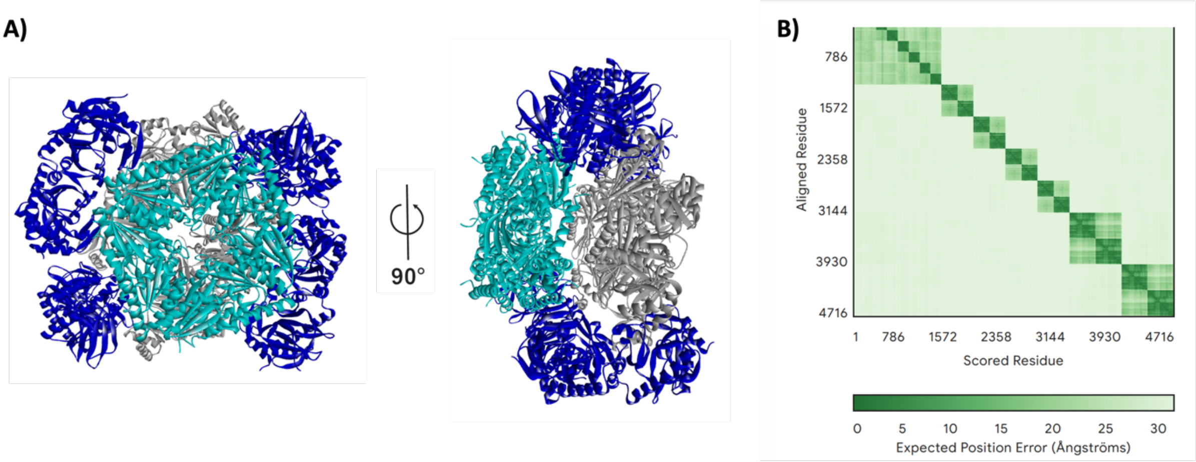


**Figure S4**. AlphaFold3 predicted structure of the PurK/PurE/PurC complex. A) Structural model of the PurK/PurE/PurC complex predicted by AlphaFold3 using the 3D-structure available in the PDB (1B6S, 1D7A and 2GQS, respectively). B) Predicted Alignment Error (PAE) plot, illustrating the expected positional uncertainty for each residue, with lower values indicating higher confidence in relative positioning.

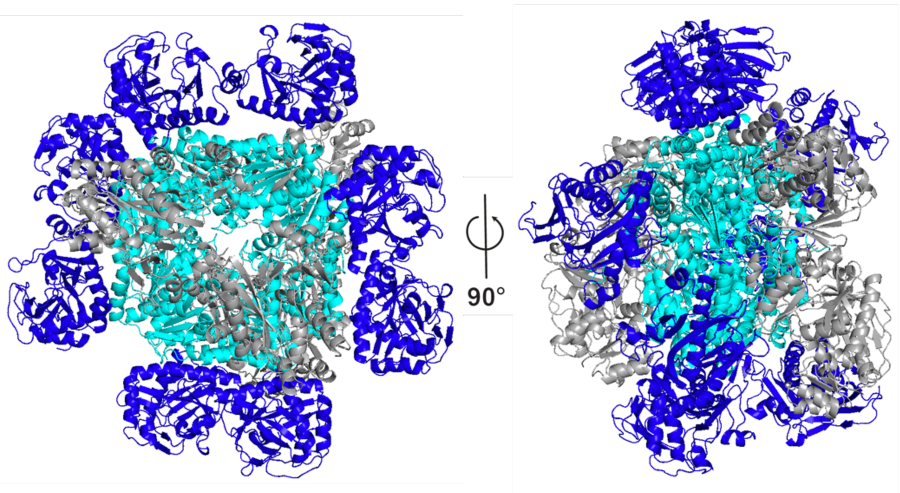


**Figure S5**. Reconstituted complex of PurK, PurE and PurC from *P. aeruginosa*. Same color code as in Fig. 6.

**TABLES**

**Table S1: full names of the enzymes involved in each step of the DNPNB pathway represented in Supplementary Figure 1.**

| | | Abbreviation/acronym | |
|---|---|---|---|
| **Step** | **Name of enzyme/domain and EC codes** | **Mammals** | ***E. coli*** |
| 1 | PRPP amidotransferase (2.4.2.14) | PPAT | PurF |
| 2 | Phosphoribosylglycinamide synthetase (6.4.3.13) | GARS domain of TrifGART | PurD |
| 3 | Phosphoribosylglycinamide formyltransferase (2.1.2.2) | GART domain of TrifGART | PurN |
| 3’ | Formate-dependent phosphoribosylglycinamide formyltransferase (6.3.1.21) | - | PurT |
| 4 | Phosphoribosyl formylglycinamidine synthase (6.3.5.3) | FGAMS | PurL |
| 5 | Phosphoribosylaminoimidazole synthetase (6.3.3.1) | AIRS domain of TrifGART | PurM |
| 6’ | $N^5$-carboxyaminoimidazole ribonucleotide synthase (6.3.4.18) | - | PurK |
| 6 | $N^5$-carboxyaminoimidazole ribonucleotide mutase (5.4.99.18) | - | PurE |
| | Phosphoribosyl aminoimidazole carboxylase (4.1.1.21) | CAIRS domain of PAICS | - |
| 7 | Phosphoribosyl aminoimidazole succinocarboxamide synthetase (6.3.2.6) | SAICARS domain of PAICS | PurC |
| 8 | Adenylosuccinate lyase (4.3.2.2) | ADSL | PurB |
| 9 | 5-aminoimidazole-4-carboxamide ribonucleotide formyltransferase (2.1.2.3) | AICART domain of ATIC | PurH |
| 10 | IMP cyclohydrolase (3.5.4.10) | IMPC domain of ATIC | PurH |
| 11 | Inosine 5’-monophosphate dehydrogenase (1.1.1.205) | IMPDH1, IMPDH2 | GuaB |
| 11’ | Adenylosuccinate synthase (6.3.4.4) | ADSS1, ADSS2 | PurA |
| 12 | Guanosine 5’-monophosphate synthase (6.3.5.2) | GMPS | GuaA |
| 12’ | Adenylosuccinate lyase (4.3.2.2) | ADSL | PurB |

**Table S2: donor vectors used in this study**

| ***Plasmid*** | ***Description*** | ***Source*** |
|---|---|---|
| pDONR201 | Gateway entry vector for inserting an attB-flanked gene | Thermofisher Scientific (Laboratory collection) |
| pTwist-ENTR | Gateway entry vector | Twist bioscience |
| pTwist -*purApa* | pTwist-ENTR harbouring the ORF of *purA* of *E. coli* | Twist bioscience |
| pTwist -*purBpa* | pTwist-ENTR harbouring the ORF of *purB* of *P. aeruginosa* | Twist bioscience |
| pTwist -*purCpa* | pTwist-ENTR harbouring the ORF of *purC* of *P. aeruginosa* | Twist bioscience |
| pTwist-*purDpa* | pTwist-ENTR harbouring the ORF of *purD* of *P. aeruginosa* | Twist bioscience |
| pDONR201-*purEpa* | pDONR201 harbouring the ORF of *purE* of *P. aeruginosa* | This work |
| pTwist -*purFpa* | pTwist-ENTR harbouring the ORF of *purF* of *P. aeruginosa* | Twist bioscience |
| pTwist -*purHpa* | pTwist-ENTR harbouring the ORF of *purH* of *P. aeruginosa* | Twist bioscience |
| pDONR201-*purKpa* | pDONR201 harbouring the ORF of *purK* of *P. aeruginosa* | [1] |
| pTwist -*purLpa* | pTwist-ENTR harbouring the ORF of *purL* of *P. aeruginosa* | Twist bioscience |
| pTwist -*purMpa* | pTwist-ENTR harbouring the ORF of *purM* of *P. aeruginosa* | Twist bioscience |
| pTwist -*purNpa* | pTwist-ENTR harbouring the ORF of *purN* of *P. aeruginosa* | Twist bioscience |
| pTwist -*purTpa* | pTwist-ENTR harbouring the ORF of *purT* of *P. aeruginosa* | Twist bioscience |
| pDONR201-*guaBpa* | pDONR201 harbouring the ORF of *guaB* of *P. aeruginosa* | This work |
| pDONR201-*ftsA* | pDONR201 harbouring the ORF of *ftsA* of *E. coli* | [2] |

**Table S3: BACTH vectors constructed and used in this study**

| *Plasmid* | *Description* | *Source or reference* |
|---|---|---|
| *pDST25-DEST* | BACTH destination vector for cloning ORF of interest at the C-terminus of T25; p15 *ori*, $Spc^r$ | [3] |
| *pUT18C-DEST* | BACTH destination vector for cloning ORF of interest at the C-terminus of T18; ColE1 *ori*, $Amp^r$ | [3] |
| pDST25-*purApa* | pST25-DEST harbouring *purA* of *P. aeruginosa* | This work |
| pDST25-*purBpa* | pST25-DEST harbouring *purB* of *P. aeruginosa* | This work |
| pDST25-*purCpa* | pST25-DEST harbouring *purC* of *P. aeruginosa* | This work |
| pDST25-*purDpa* | pST25-DEST harbouring *purD* of *P. aeruginosa* | This work |
| pDST25-*purEpa* | pST25-DEST harbouring *purE* of *P. aeruginosa* | This work |
| pDST25-*purFpa* | pST25-DEST harbouring *purF* of *P. aeruginosa* | This work |
| pDST25-*purHpa* | pST25-DEST harbouring *purH* of *P. aeruginosa* | This work |
| pDST25-*purKpa* | pST25-DEST harbouring *purK* of *P. aeruginosa* | [1] |
| pDST25-*purLpa* | pST25-DEST harbouring *purL* of *P. aeruginosa* | This work |
| pDST25-*purMpa* | pST25-DEST harbouring *purM* of *P. aeruginosa* | This work |
| pDST25-*purNpa* | pST25-DEST harbouring *purN* of *P. aeruginosa* | This work |
| pDST25-*purTpa* | pST25-DEST harbouring *purT* of *P. aeruginosa* | This work |

| | | |
|---|---|---|
| pDST25-*guaBpa* | pST25-DEST harbouring *guaB* of *P. aeruginosa* | This work |
| pKT25-*ftsA* | pKT25-DEST harbouring *ftsA* of *E. coli* | [2] |
| pKT25-zip | pKT25 encoding Zip, the leucine zipper region of the yeast protein GCN4 | [4] |
| pUTC18C-*purApa* | pUT18C-DEST harbouring *purA* of *P. aeruginosa* | This work |
| pUT18C-*purBpa* | pUT18C-DEST harbouring *purB* of *P. aeruginosa* | This work |
| pUT18C-*purCpa* | pUT18C-DEST harbouring *purC* of *P. aeruginosa* | This work |
| pUT18C-*purDpa* | pUT18C-DEST harbouring *purD* of *P. aeruginosa* | This work |
| pUT18C-*purEpa* | pUT18C-DEST harbouring *purE* of *P. aeruginosa* | This work |
| pUT18C-*purFpa* | pUT18C-DEST harbouring *purF* of *P. aeruginosa* | This work |
| pUT18C-*purHpa* | pUT18C-DEST harbouring *purH* of *P. aeruginosa* | This work |
| pUT18C-*purKpa* | pUT18C-DEST harbouring *purK* of *P. aeruginosa* | [1] |
| pUT18C-*purLpa* | pUT18C-DEST harbouring *purL* of *P. aeruginosa* | This work |
| pUT18C-*purMpa* | pUT18C-DEST harbouring *purM* of *P. aeruginosa* | This work |
| pUT18C-*purNpa* | pUT18C-DEST harbouring *purN* of *P. aeruginosa* | This work |
| pUT18C-*purTpa* | pUT18C-DEST harbouring *purT* of *P. aeruginosa* | This work |
| pUT18C-*guaBpa* | pUT18C-DEST harbouring *guaB* of *P. aeruginosa* | This work |
| pUT18C-*ftsA* | pUT18C harbouring *ftsA* of *E. coli* | [2] |

| pUT18C-zip | pUT18C encoding Zip, the leucine zipper region of the yeast protein GCN4 | [4] |
|---|---|---|

**Table S4: Adjusted p-values (a.p-values) calculated from the means and standard deviations of β-galactosidase activity measurements in BACTH assays, performed on eight colonies for each pair of T25 (row) and T18 (column) fusion proteins.**

| | PurA | PurB | PurC | PurD | PurE | PurF | PurH | PurK | PurL | PurM | PurN | PurT | IMPDH | ftsA |
|---|---|---|---|---|---|---|---|---|---|---|---|---|---|---|
| PurA | 3.57E-14 | 0.88148148 | 0.88148148 | 0.38233809 | 0.88148148 | 0.88148148 | 0.88148148 | 0.03556688 | 0.88148148 | 0.88148148 | 0.00052252 | 0.88148148 | 0.88148148 | 0.88148148 |
| PurB | 0.76502422 | 3.57E-14 | 0.6070319 | 0.04060945 | 0.01413686 | 0.88148148 | 0.0099366 | 3.57E-14 | 0.88148148 | 0.0101024 | 0.88148148 | 0.03139063 | 0.01841385 | 0.88148148 |
| PurC | 0.88148148 | 3.62E-05 | 4.25E-12 | 0.88148148 | 0.88148148 | 0.88148148 | 0.63368074 | 6.17E-06 | 0.88148148 | 0.88148148 | 0.88148148 | 0.38233809 | 0.88148148 | 0.88148148 |
| PurD | 0.88148148 | 0.88148148 | 0.88148148 | 0.88148148 | 0.33818411 | 0.00034979 | 0.88148148 | 0.16469568 | 0.88148148 | 0.88148148 | 0.88148148 | 0.88148148 | 0.05372834 | 0.88148148 |
| PurE | 1.71E-08 | 4.52E-09 | 0.88148148 | 0.50404926 | 3.57E-14 | 0.80744169 | 1.20E-11 | 1.06E-05 | 0.88148148 | 1.59E-07 | 0.15158361 | 3.65E-12 | 0.63368074 | 0.88148148 |
| PurF | 0.88148148 | 0.88148148 | 0.00314496 | 0.04934308 | 0.00782674 | 0.88148148 | 0.0010021 | 0.88148148 | 0.21756776 | 0.09380523 | 0.88148148 | 0.88148148 | 0.11532438 | 0.88148148 |
| PurH | 0.00979406 | 5.99E-14 | 2.09E-10 | 0.88148148 | 6.31E-14 | 0.63368074 | 3.57E-14 | 0.88148148 | 0.88148148 | 0.02579022 | 0.88148148 | 0.88148148 | 0.00526315 | 0.88148148 |
| PurK | 0.01699269 | 0.88148148 | 0.88148148 | 0.88148148 | 0.88148148 | 0.88148148 | 0.88148148 | 2.09E-10 | 0.88148148 | 0.88148148 | 0.88148148 | 0.88148148 | 0.01854607 | 0.88148148 |
| PurL | 0.88148148 | 0.88148148 | 0.88148148 | 0.88148148 | 0.88148148 | 3.49E-05 | 0.88148148 | 0.88148148 | 0.88148148 | 0.88148148 | 0.88148148 | 0.88148148 | 0.88148148 | 0.88148148 |
| PurM | 0.88148148 | 0.88148148 | 0.88148148 | 0.88148148 | 0.88148148 | 6.31E-14 | 0.88148148 | 0.88148148 | 0.88148148 | 4.70E-09 | 0.88148148 | 0.88148148 | 0.88148148 | 0.88148148 |
| PurN | 0.88148148 | 0.88148148 | 0.88148148 | 0.88148148 | 0.88148148 | 0.88148148 | 0.88148148 | 0.88148148 | 0.88148148 | 0.34909133 | 0.88148148 | 0.01367607 | 0.52243586 | 0.88148148 |
| PurT | 0.45352417 | 0.88148148 | 0.88148148 | 0.88148148 | 9.78E-07 | 0.16469568 | 0.88148148 | 0.88148148 | 0.88148148 | 0.88148148 | 0.88148148 | 0.11532438 | 0.88148148 | 0.88148148 |
| IMPDH | 0.88148148 | 0.56957851 | 0.88148148 | 0.88148148 | 0.88148148 | 0.88148148 | 0.88148148 | 0.88148148 | 0.88148148 | 0.88148148 | 0.88148148 | 0.88148148 | 7.29E-08 | 0.88148148 |
